# Gene-Family Encoding Boosts Domain-Adapted Single-Cell Language Models

**DOI:** 10.1101/2025.09.16.676672

**Authors:** Haoran Ma, Chang Xu, Shamaine Wei Ting Ho, Joseph J Zhao, Yunqiang Chu, Angie Lay Keng Tan, Raghav Sundar, Patrick Tan

## Abstract

Transformer-based single-cell foundation models often rely on ranked-gene (RG) sequences where genes, ranked by expression, are often not functionally related, weakening next-token learning and the structure of learned embeddings. Here, we introduce gene-family (GF) encoding, where expressed genes are grouped into functionally defined families, and ranking performed within each family. Using 100,000 gastric-cancer (GC) cells, we domain-adapted 8-billion-parameter Llama and Qwen backbones with either RG or GF sentences and benchmarked zero-shot performance on embedding-based and generative tasks. GF models outperformed RG models across both tasks and backbones. We scaled GF-Llama to 1.3 million GC cells to obtain GF-Llama-GC and applied it to two applications: resolving fine-grained cellular heterogeneity and discovering cell populations associated with disease progression. GF-Llama-GC revealed immune-cell subclusters not resolved by standard expression-based analyses. Applying *in-silico* cell removal/transplantation on a chemotherapy responder/non-responder scRNA-seq dataset, GF-Llama-GC highlighted not only epithelial cells but also neutrophils as key cells associated with chemotherapy response.

## Introduction

Artificial-intelligence (AI) driven transformer models are being increasingly applied to the emerging field of single-cell biology. Recent examples such as Geneformer [1], scGPT [2] and scFoundation [3] treat genes as tokens and learn general representations, which can then be adapted to various tasks including cell-type annotation, batch correction, data integration, *in silico* gene perturbation and predicting cell–cell interactions. However, while flagship language models [4–6] now span tens to hundreds of billions of parameters, current single-cell foundation models remain in the hundreds of millions range (Geneformer ≈ 0.3 B; scGPT ≈ 0.05 B; scFoundation ≈ 0.1 B). The latter’s limited capacity restricts what these models learn, as scaling-law research has shown that larger networks frequently achieve decreased pre-training loss [7] and transfer more effectively after task-specific adaptation [8, 9]. Consistent with this limitation, existing single-cell models have been shown to underperform on demanding downstream tasks such as *in silico* perturbation prediction [10, 11].

Scaling single-cell models from hundreds of millions to billions of parameters is therefore attractive but costly, as training *de novo* models of that size requires substantial computational and financial resources [12, 13]. Moreover, single-cell RNA sequencing (scRNA-seq) data sets, when taken in isolation, lack key components of biological information, such as protein-protein interactions, regulatory influences from non-expressed cis-regulatory elements, and evolutionary conservation. To incorporate these other data types, domain-adaptive pre-training offers an attractive alternative, since many contemporary large, general-purpose language models already encode diverse biomedical knowledge including gene symbols after being exposed to trillions of tokens from public databases and the existing literature. Adapting such general-purpose models with single-cell transcript information is also likely to be far less expensive than training from scratch and can also inject cell-state–specific signals into a knowledge-rich backbone. Indeed, a recent study showed that continued pre-training of the Gemma-2 family on scRNA-seq data reported consistent performance gains as parameter count increased, supporting scaling-law predictions [14].

However, to realize the potential of domain-adaptive pre-training, single-cell foundation models still face two major challenges. First, there are model–data mismatches – specifically, most current single-cell models adapt the transformer architecture, which infers the probability of the next token given the entire preceding context. For single-cell RNA-seq, the tokens are often gene names whose order is derived from expression values or other heuristics [1–3]. However, because neighbouring genes in such sequences can be functionally unrelated (e.g., mitochondrial *MT-ND4* can be followed by a cell-type marker such as *EPCAM*), true biological relationships can be sparse and noisy. Recent benchmarking studies report that transformers trained on ranked-gene sequences capture broad cell states but struggle with fine-grained functional relationships [15]. The second challenge is the breadth of single-cell data. Model capacity scales with both parameter count and training diversity [16, 17]. The latest CELLxGENE release (v2025-08-01) aggregates 123 million cells across 1,869 datasets and > 1,000 annotated cell types [18]. However, even within this large dataset, coverage remains uneven: gastric cancer (GC) samples, for instance, comprise ∼294,000 cells from a single cohort, offering limited variation in genotype, stage and micro-environment. Such narrow diversity restricts a model’s ability to generalise to unseen diseases or rare perturbations. Together, these architectural and data constraints inevitably limit performance on demanding tasks (eg *in silico* gene-perturbation prediction, cross-disease transfer learning). Addressing these limitations thus requires representations such as set-based encodings that can expose non-sequential gene relationships and which can integrate with broader and more balanced training corpora.

In this study, we introduce a new single-cell representation - gene-family (GF) encoding. In this representation, we first grouped the transcriptome into 29 manually curated gene families and then ranked genes by their expression levels within each family to yield function-focused token sequences. Focusing on GC, we assembled 1.3 million cells from 10 public studies and one in-house cohort with matched clinical data. We then performed domain-adaptive pre-training using two 8-billion-parameter language models Llama-3.1-8B and Qwen-3-8B on the GF cell sentences. On three knowledge benchmarks using embeddings (gene co-expression network inference, gene sets clustering and intra/inter group similarity) and two generative tasks (next-token prediction and whole cell stimulation), the GF models consistently outperformed rank-based counterparts. In terms of biological insights, embeddings from the GF models identified heterogeneous T-cell states in a GC cohort where conventional algorithms failed to find, and also resolved cellular heterogeneity in cancers beyond GC, such as ovarian peritoneal metastasis. Finally, using our single-cell GF model, we developed a novel system which identifies key cells driving state changes at the patient level. Applied to chemotherapy responder versus non-responder samples, the GF method highlighted not only epithelial cells, but also neutrophils, as key cell types influencing chemotherapy response in GC.

## Results

### Overview of data preparation and model training

Public resources such as CELLxGENE aggregate scRNA-seq datasets across diseases, but coverage for any single cancer is uneven and often concentrated in a small number of cohorts with limited clinical annotation. For example, in GC, the publicly available cells in CELLxGENE are only ∼294,000 from a single cohort, which is insufficient for training high-capacity single-cell language models. We therefore created a new GC single-cell atlas dataset, combining ten public GC datasets and one in-house cohort (**Supplementary Table 1**). After performing manually processed quality control, normalization, clustering and refined cell annotation (see Methods), the resulting GC atlas comprises 1,313,051 cells distributed across 17 annotated cell types (**Figure 1A, Supplementary** Figure 1A). Most cell lineages show comparable abundance across clinical subtypes, whereas macrophages were enriched in stage-IV tumours (**Figure 1B**). As an alternative to conventional ranked-gene (RG) representations, we developed GF encoding (**Figure 1C**). In GF encoding, genes are first ranked by their expression within each cell. Second, the ranked list is then partitioned into 29 curated gene families, 15 reflecting general cellular programmes and 14 enriched in tumours (**Supplementary Table 2**). Third, one short sub-sentence is created for every family, and all sub-sentences are concatenated to form the final cell sentence (See **Methods**). Gene-overlap analysis shows that families tagged as “tumour” share very few genes with normal families; but exhibit overlap with related families such as *cell-death* and *oncogene* clusters (**Figure 1D**).

**Figure 1.**
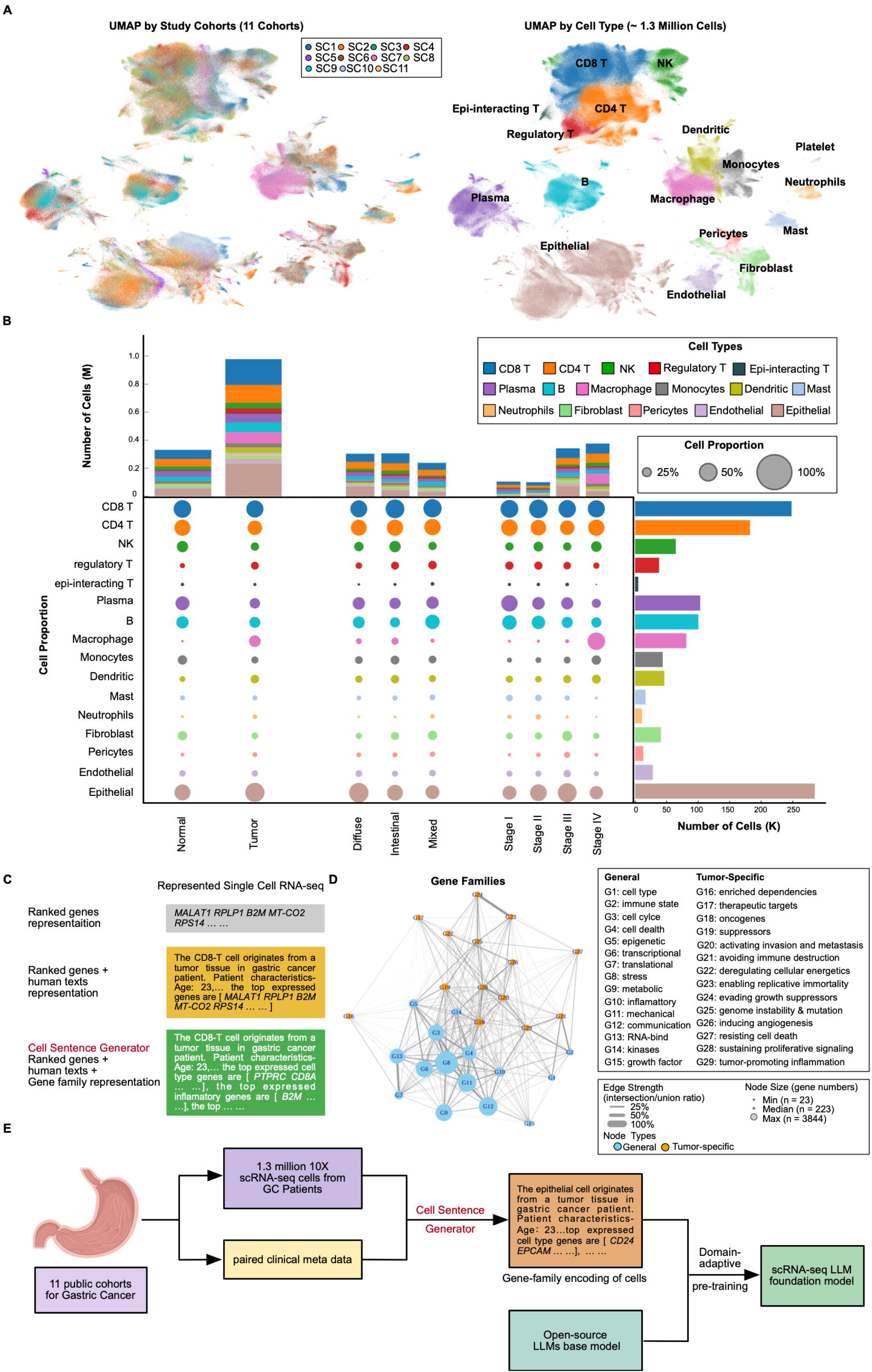
**Overview of data and model** A. UMAP of the 1.3 million cells from 11 public gastric cancer cohorts, colored either by dataset (left) or by annotated cell types (right). The overall dataset contains 1,313,051 cells and 36,017 genes after quality control, and was annotated to 17 cell types. B. Cell type compositions of integrated GC scRNA-seq data. The middle panel (bubble plot) shows the cell subclusters (rows) by: adjacent normal samples vs tumor samples, by Lauren subtypes (diffuse vs. intestinal vs. mixed), and by tumor stages (I-IV). The size of the circle represents the cell proportion of each specific cell type. The circles are colored by annotated cell types. The stacked bar graph at the top shows the number of cells for each category. The histogram on the right shows the absolute cell numbers for each cell type. Platelet cell was excluded because of the low cell number. C. Schematic depiction of 3 methods of scRNA-seq representation. The single-cell sentence in the grey box is generated using a conventional expression rank-based encoding method. The single-cell sentence in the yellow box added clinical information based on the first method. The single-cell sentence in the green box is generated using our defined gene-family encoding method. D. Gene family network plot of the defined 29 gene families. Each node represents a gene family, the size of the node shows the number of genes in this category. Node was colored by whether the gene family belongs to general or tumor-specific gene families. The edge represents the gene intersection-union ratio between two gene families, and the width was normalized across all edges. The text “G1” to “G29” on top of the node represent gene family 1 to gene family 29. E. Workflow depiction for adaptive pretraining on public open-source large language models using our GC cell sentences. The “cell sentence generator” here specifically refers to the gene-family encoding method.

To construct the GC single-cell foundation model, we then generated 1.32 million cell sentences from the curated GC atlas (**Figure 1E**). Each sentence was produced using the GF encoding based generator (**Methods**). To provide comparisons, we also performed RG encoding. We then implemented domain-adaptive pre-training using two 8-billion-parameter backbones Llama3.1-8B and Qwen3-8B on the GF and RG sentences to obtain single-cell language models. Using both RG and GF encoders, we first created 100,000 training sentences and trained four pilot models including GF-Llama, RG-Llama, GF-Qwen, and RG-Qwen under identical hyper-parameters. We performed systematic benchmarking on these 4 models (see sections below), and finally retrained the best backbone on the full 1.3 million sentences to produce the final GF-Llama-GC foundation model.

### Gene-family encodings strengthen biological structure in single-cell embeddings

Previous work has shown that in domain-adaptive pre-training, large mismatches in token distribution or syntax between the pre-training corpus and the backbone model’s original training data can impair convergence and downstream performance [19, 20]. Single-cell “sentences”, which mix natural-language metadata with gene symbols, represent a non-intuitive and hybrid linguistic structure that may reduce adaptation efficiency when applied to general-purpose language models. To assess whether GF encoding produces single-cell sentences that are closer to natural-language structure than RG encoding, we used the Gemini-2.0-flash model to assign a “readability score” to 100 randomly sampled sentences from each representation (see **Methods**). GF-encoded sentences achieved a mean readability score 28.87% higher than RG-encoded sentences (**Supplementary** Figure 2A). We then performed domain-adaptive pre-training for one epoch on 100,000 cells using two public 8-billion-parameter backbones Llama3.1-8B Base and Qwen3-8B with either GF- or RG-encoded sentences. In the Llama backbone, GF-encoded training reduced final losses by 0.52 compared with RG encoding (**Supplementary** Figure 2B), indicating faster convergence during adaptation.

Next, we evaluated if models trained with different single-cell representations and backbone architectures GF-Llama (gene-family, Llama-3.1-8B), RG-Llama (ranked-gene, Llama3.1-8B), GF-Qwen (gene-family, Qwen3-8B), and RG-Qwen (ranked-gene, Qwen3-8B) captured biologically meaningful relationships during domain-adaptive pre-training (**Figure 2A**). For comparison, we also included three reference models: Geneformer (a representative single-cell foundation model using RG encoding without domain-adaptive pre-training), the unadapted Llama3.1-8B, and the unadapted Qwen3-8B. Unlike previous benchmarking studies, which have mainly assessed downstream tasks that require further fine-tuning such as cell-type prediction, batch-effect removal and *in silico* perturbation [1–3], we focused on the zero-shot setting. Specifically, we assessed each model’s ability to encode biological structure and relationships directly from pre-training, using two complementary evaluation streams: embedding-based benchmarks (at the gene level and the gene-program level) and generation benchmarks (at the token level and the sentence level) (**Figure 2A**).

**Figure 2.**
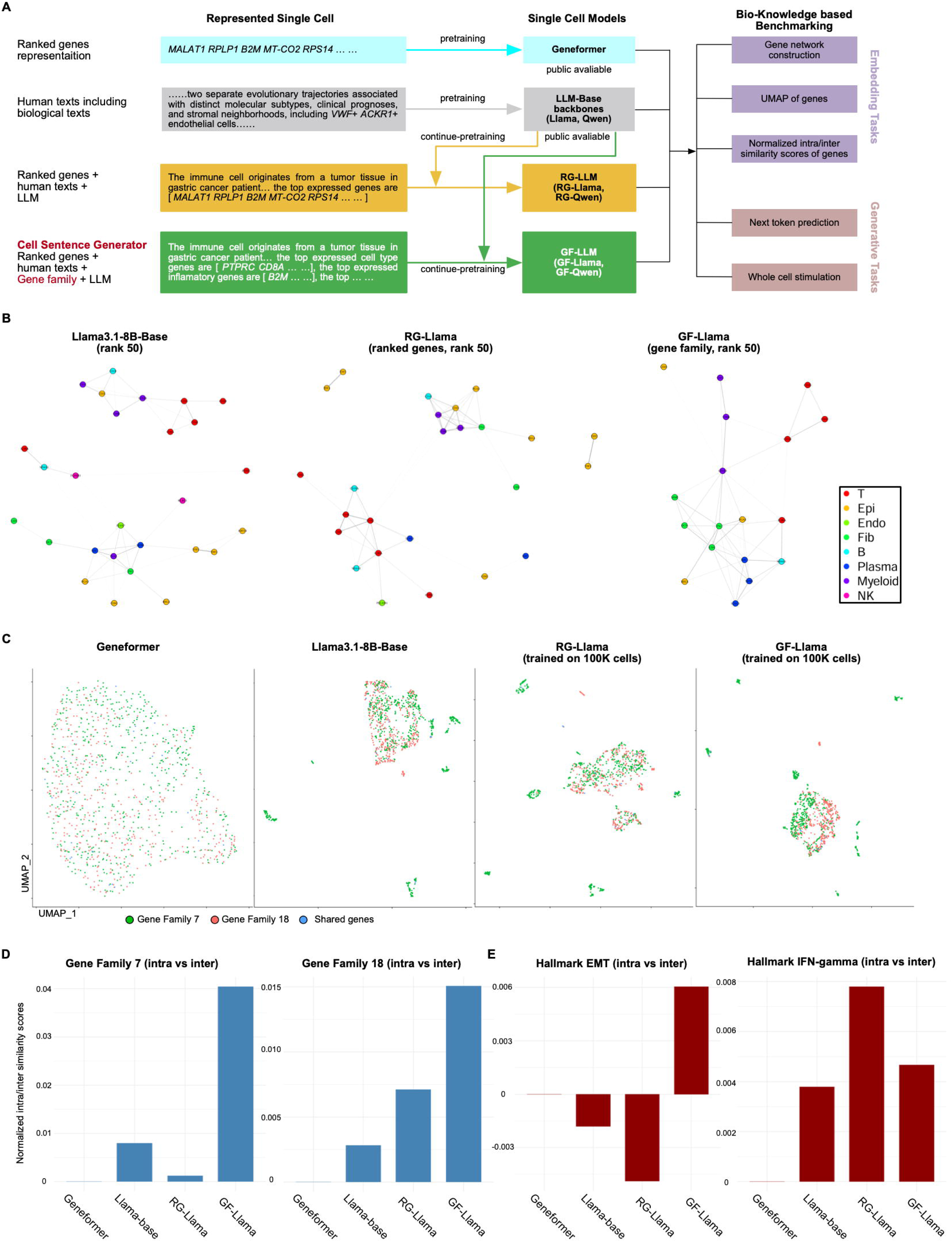
**Benchmarking using embeddings of models** A. Workflow depiction for benchmarking different models trained by various single-cell representation methods and backbones. Geneformer was selected as a representative RG-encoding trained model which only used scRNA-seq data. Open-source LLMs including Llama3.1-8B-base and Qwen3-8B were selected as representative LLM backbones. We also perform domain-adaptive pretraining on the 2 backbones using either GF- or RG-encoding generated cell sentences, yielding another 4 models (GF-Llama, RG-Llama, GF-Qwen, RG-Qwen). These models were benchmarked on a series of biological tasks including both the embedding-based tasks and generative tasks. B. Gene correlation network plot defined using KNN and cosine similarities of cell-type-specific gene embeddings for Llama models. Each dot represents a gene, and the dot was colored by cell types. The link between two dots represents that these two genes have a high cosine similarity ranked top 50 among all gene pairs. C. UMAP projections of gene embeddings from a mixture of defined Gene Family 7 (protein translation-related genes) and Gene Family 18 (oncogene-related genes) for Geneformer and models using the Llama backbone. Each dot represents a gene, and the dot was colored by different gene families. D. Barplots showing the defined normalized intra/inter cosine similarity scores on Gene Family 7, Gene Family 18 and 2 unseen Hallmark pathways (EMT and IFN-ψ) that are not defined in gene families in Llama models. Positive values show the model can distinguish among different gene programs/pathways, and the magnitude of the values represents how well it can separate gene programs/pathways.

For the gene-level embedding zero-shot benchmark, we evaluated whether each model could distinguish biologically-related from unrelated genes using the cosine similarity of their embedding vectors (**Supplementary** Figure 2C). Three gene sets were defined: (i) *null***-** randomly selected genes (5000 random pairs) to estimate the background similarity distribution; (ii) *positive control***-** ribosomal protein large subunit (RPL) genes, which share highly similar biological functions and are housekeeping genes; and (iii) *negative control***-** genes drawn from distinct immune programmes (inhibitory checkpoints, stimulatory checkpoints, chemokines, cytokines and MHC). In the null set, Qwen3-8B (RG, GF and unadapt) and Geneformer exhibited high mean similarity (0.9868-0.9870) across random pairs, suggesting limited discriminative capacity. In contrast, Llama3.1-8B (RG, GF and unadapt) showed lower background similarity (0.8519-0.9031), allowing clearer separation between gene sets. When comparing GF and RG encodings, only GF-Llama captured correct directionalities (positive control > null > negative control), with statistically distinct similarity distributions for both positive and negative controls relative to null (Kolmogorov–Smirnov test, p = 0.01). All other models failed to capture the correct directionality.

We then assessed whether embeddings could recover cell-type-specific gene relationships. Marker genes for epithelial cells, T cells, NK cells, B cells, plasma cells, myeloid cells, fibroblasts and endothelial cells were curated (**Supplementary Table 2**), and gene co-expression networks were inferred from pairwise embedding correlations (neighbours = 10, rank = 50) (**Figure 2B, Supplementary** Figure 2D). Geneformer and the unadapted backbones produced mixed networks with cell-type markers interspersed. RG-Llama, RG-Qwen and GF-Qwen mainly grouped genes with identical prefixes but are otherwise biologically unrelated (e.g., *CD3E* and *CD24*), consistent with subword-tokenisation artefacts. Only GF-Llama recovered coherent, cell-type-specific clusters for all five lineages, indicating that gene-family encoding enhances the model’s ability to capture biologically meaningful gene-gene associations.

At the gene-program level, we then assessed whether embeddings could distinguish functionally distinct gene sets. As an initial test, we selected two previously defined gene families: Gene Family 7 (protein translation-related genes) and Gene Family 18 (oncogene-related genes) from **Supplementary Table 2**, and projected their embeddings from each model into two dimensions using UMAP (see Methods).

For both Llama3.1-8B and Qwen3-8B backbones, models trained with GF-encoded cells produced visually distinct clusters for the two families. In contrast, Geneformer, the unadapted backbones, and models trained with RG-encoded cells showed substantial overlap between families (**Figure 2C, Supplementary** Figure 2E). To further quantify this separation, we calculated a normalised intra/inter-program similarity ratio (see **Methods**), where values > 0 indicate higher within-program similarity relative to between-program similarity. Using Gene Families 7 and 18, GF-Llama achieved the highest ratio among all models (**Figure 2D, Supplementary** Figure 2F). We then evaluated the generalisability of this finding using two Hallmark pathways not included in the predefined families, including epithelial–mesenchymal transition (EMT) and IFN-γ signalling. Again, GF-Llama produced high similarity ratios for both pathways, whereas other models yielded low or negative ratios (intra-program similarity below inter-program similarity), especially for EMT (**Figure 2E, Supplementary** Figure 2G). These results indicate that GF encoding enhances the model’s ability to separate biologically distinct gene programs, even when they are unseen during training. Overall, our zero-shot benchmarking demonstrates that GF encoding consistently improves the ability of large language model backbones to capture biologically meaningful structure from single-cell data without additional fine-tuning.

### Gene-family encoding improves generative performance in single-cell foundation models

The transformer architecture allows autoregressive generation, an application absent from many earlier single-cell foundation models [1–3]. To test whether GF encoding also improves generative performance, we conducted a gene-level next-token prediction task. Using two 8-billion-parameter backbones (Llama3.1-8B and Qwen3-8B), we compared models trained on GF-versus RG-encoded sentences. Each model was prompted with 1,000 cell descriptions consisting of natural-language metadata and two cell–type–specific marker genes (see **Methods**). For each prompt, we recorded the top five predicted next gene tokens and their associated likelihoods. In GF-Llama, the likelihood distribution was strongly concentrated indicating high prediction confidence, with the top prediction accounting for ∼40% of total probability mass, and the top five collectively exceeding 80% (**Figure 3A; Supplementary** Figure 3A). By contrast, RG-Llama, GF-Qwen and RG-Qwen showed more uniform distributions across candidate tokens, indicating decreased prediction certainty. These results support our hypothesis that grouping genes into functional families provides contextual cues that make the next gene more predictable—for example, “*CD3*” is more probable when preceded by other immune cell–related genes, thereby reducing uncertainty in generative tasks.

**Figure 3.**
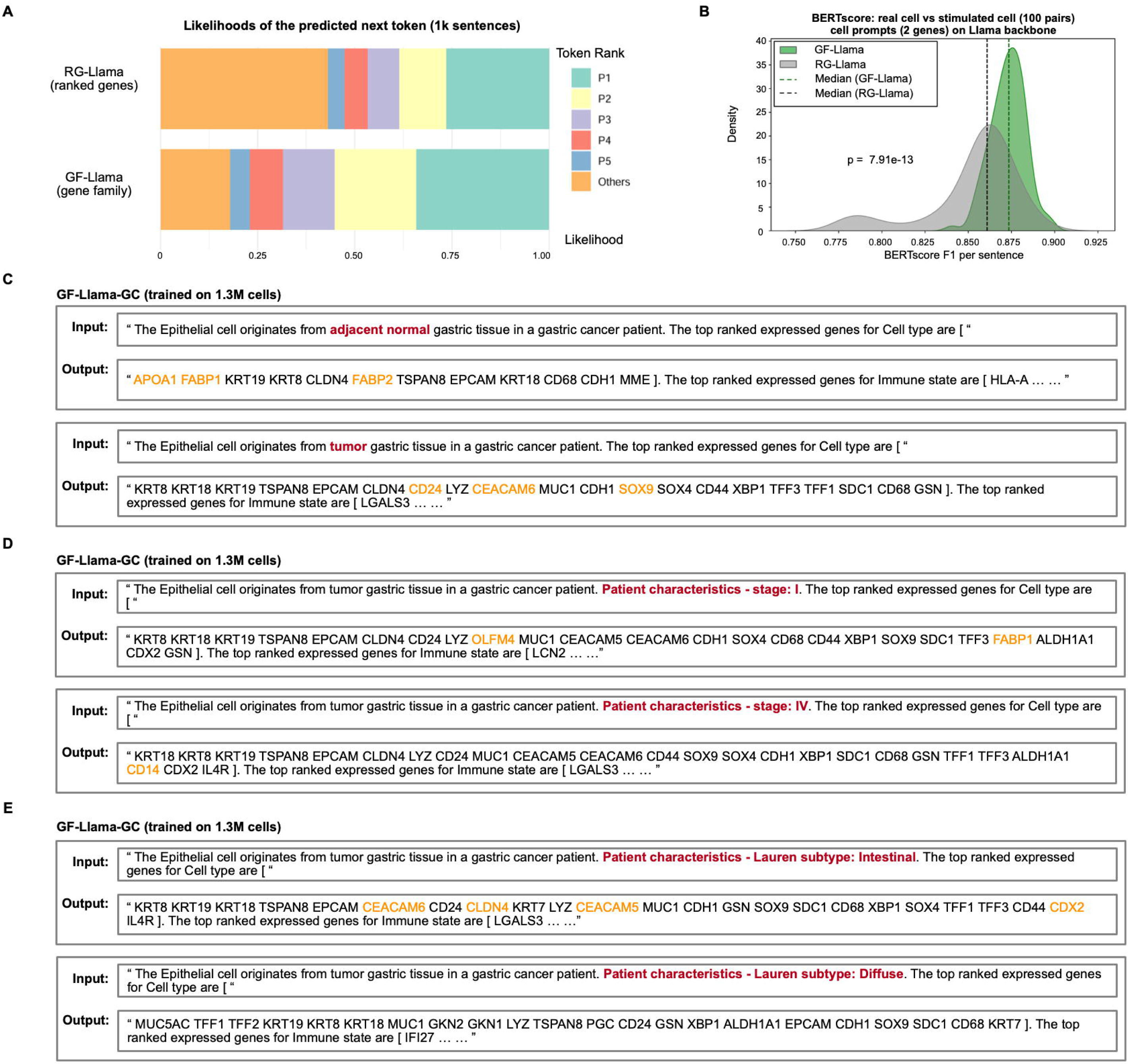
**Benchmarking on generative ability** A. Barplots showing the distribution of the likelihood of the next predicted token from GF-Llama and RG-Llama. The likelihood of the top 5 candidate tokens and the rest was colored. The result takes the mean value from 1,000 sentences. B. Density plot of the BERTscore F1 values for GF-Llama and RG-Llama between 100 pairs of stimulated cell sentences (with 2 hint genes) and reference sentences. The dotted line indicates the median value of each model. Colors represent the F1 values from different models. P-value is 7.91e^-13^ by the two-sided Mann–Whitney U test. C. Representative examples of stimulated cell sentences under adjacent normal versus tumor settings using GF-Llama-GC. Key cell-type-specific markers were highlighted in yellow. D. Representative examples of stimulated cell sentences under adjacent normal versus tumor settings using GF-Llama-GC. Key cell-type-specific markers were highlighted in yellow. E. Representative examples of stimulated cell sentences under adjacent normal versus tumor settings using GF-Llama-GC. Key cell-type-specific markers were highlighted in yellow.

Next, at the whole-cell level, we assessed each model’s ability to generate complete cell sentences. Using the same prompts as in the next-token prediction task (cell metadata followed by two cell–type–specific marker genes from 100 randomly selected real cells), we allowed the models to continue generating the remaining tokens until they reached 500 genes. The generated sentences were compared with corresponding real-life cells using the BERT score [21], which measures semantic similarity between token sequences. In both Llama3.1-8B and Qwen3-8B backbones, models trained with GF-encoding achieved significantly higher BERT scores than those trained with RG-encoding (**Figure 3B; Supplementary** Figure 3B), indicating that GF-encoded models produce cell sequences more similar to real cells. We then applied a stricter evaluation, using only the cell metadata as prompts without providing the first two genes as hints. Under this setting, GF-encoded models again outperformed RG-encoded models for both backbones (**Supplementary** Figure 3C**-D**), with performance metrics very close to those using hint genes. This result demonstrates that gene-family encoding enhances generative performance even with minimal input context.

Beyond evaluating the generative ability of the mini-models trained on 100,000 cells for one epoch, we then developed and tested GF-Llama-GC, a scaled-up version of GF-Llama trained on all 1.3 million cells for three epochs. Previous single-cell generation studies have typically conditioned on cell type alone [14]. Here, we examined whether the model could generate cell sentences when provided with additional clinical contexts, such as histological subtypes or disease stages. For example, when prompted with descriptions of the same epithelial cell type but from different sample types such as tumour versus adjacent normal tissue, the generated cell sentences exhibited distinct marker profiles (**Figure 3C**). In simulated adjacent-normal epithelial cells, top-ranked markers included *APOA1* [22], *FABP1* [22] and *FABP2* [23], which are intestinal metaplasia cell markers at a pre-cancer stage. In contrast, simulated tumour epithelial cells were enriched for tumour-associated markers such as *CD24* [24], *CEACAM6* [25], and *SOX9* [26], in line with known GC biology. Similarly, when generating tumour epithelial cells from different pathological stages (**Figure 3D**), simulated tumor stage I cells expressed intestinal metaplasia– associated markers including *OLFM4* [23] and *FABP1* [23], indicative of pre- or early-stage disease. Simulated stage IV cells were enriched for markers associated with epithelial–mesenchymal transition (EMT), such as *CD14* [27]. We also evaluated performance across Lauren histological subtypes (**Figure 3E**). Simulated intestinal-type epithelial cells contained characteristic intestinal markers such as *CDX2* [28, 29], CLDN4 [30], *CEACAM5* [31] and *CEACAM6* [25], whereas these markers were absent from simulated diffuse-type epithelial cells. Together, these results demonstrate that GF-Llama-GC can generate biologically coherent single-cell sentences conditioned on diverse clinical attributes, highlighting its potential for *in silico* modelling of context-specific cell states.

Interestingly, our results also revealed under-appreciated nuances of specific backbone models on the performance of domain-adaptive pre-trained single-cell models. Qwen3 is a highly ranked open-source large language model that has outperformed Llama3.1 on several general-purpose LLM leaderboards (eg https://lmarena.ai/leaderboard, https://huggingface.co/spaces/open-llm-leaderboard/open_llm_leaderboard#/). Unexpectedly, in our single-cell foundation model benchmarks, Llama3.1 consistently achieved better performance than Qwen3 across most evaluations. We hypothesise that differences in tokenisation may be a key factor. For example, when converting gene symbols to tokens, Qwen’s tokenizer produces a median of five tokens per gene, compared with three tokens using Llama’s tokenizer (**Supplementary** Figure 3E). Because gene embeddings are computed by averaging token embeddings, a larger number of tokens per gene increases the chance of shared tokens—particularly for genes with similar prefixes (e.g., *CD3E*, *CD3D*), thereby inflating their similarity scores. This effect reduces Qwen’s ability to distinguish between biologically unrelated genes. Other possible contributing factors include differences in token vocabulary size, the proportion of English-language text in the original training data (as most gene symbols are English), and the extent of biology- and medicine-related content in the training corpus. Based on these findings, we recommend that researchers performing domain-adaptive pre-training of single-cell foundation models systematically compare multiple LLM backbones and tokenizers, rather than relying solely on general-purpose LLM leaderboard rankings, which do not reflect performance in specialised biomedical contexts.

In summary, GF encoding consistently improved the generative capacity of single-cell foundation models, enabling accurate prediction of context-relevant genes and realistic reconstruction of complete cell profiles. The scaled-up GF-Llama-GC model could generate biologically coherent single-cell representations conditioned on diverse clinical attributes, such as tissue origin, disease stage, and histological subtype, capturing known marker gene patterns.

### Gene-family–based embeddings uncover fine-grained cellular heterogeneity

Two major applications of scRNA-seq in biomedical research include the identification of rare or novel cell types/states in biological samples associated with disease outcomes [32–36], or shifts in the relative abundance and composition of more prevalent cell types that can also drive disease progression [37–40]. To investigate if GF-based single-cell foundation models can be applied to these settings and improve sensitivity compared with conventional approaches, we applied GF-Llama-GC cell embeddings to evaluate their ability to resolve cellular heterogeneity (**Figure 4A**). To validate whether embeddings derived from “cell sentences” can capture cell states and their relationships, we used an intestinal metaplasia (IM) dataset which contains well-defined gastric and intestinal epithelial lineages along with other cell types [23]. Notably, this IM dataset assembled across 18 patients, representing a precancerous rather than overtly malignant condition, was unseen during model training and was not used for fine-tuning, thus providing a stringent zero-shot generalization test of cell-state representation. In the original study using conventional scRNA-seq analytical pipelines, gene expression data was processed to capture both gastric and intestinal epithelial lineages, and their neighbourhood relationships were visualized in UMAP space. In the current approach, cells were first represented as cell sentences using GF encoding, then passed through GF-Llama-GC to obtain embeddings, which were subsequently scaled, reduced in dimensionality, and visualized with UMAP (see **Methods**). The resulting UMAP showed that cells were appropriately clustered by cell type rather than by patient batch effects (**Figure 4B**). Importantly, both gastric and intestinal epithelial lineages were correctly recovered, with neighbourhood relationships among cell groups closely matching those reported in the original study. These results indicate that GF-Llama-GC embeddings can robustly capture cell states even in unseen, non-GC datasets, importantly recapitulating findings similar to conventional scRNA-seq analysis pipelines.

**Figure 4.**
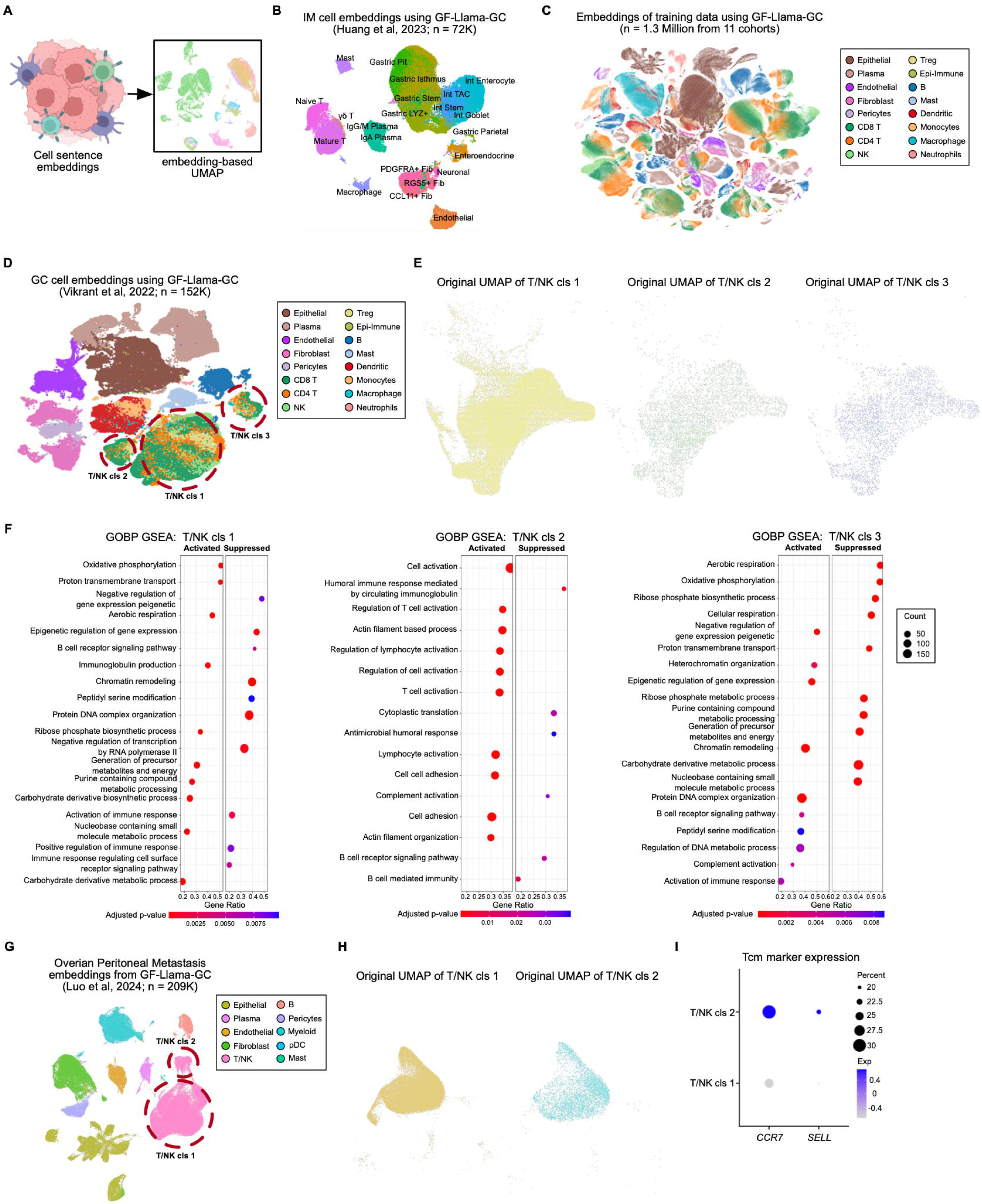
**GF embedding dissolved cell heterogeneity** A. Schematic depiction of using cell sentence embeddings to generate UMAP for visualizing possible cell heterogeneity. B. The model embedding-generated UMAP from an intestinal metaplasia dataset (Huang et al). Each dot represents a single cell. Dots are colored by annotated cell types from scRNA-seq analysis. C. The model embedding-generated UMAP from all 1.3 million training cell sentences from 11 cohorts. Each dot represents a single cell. Dots are colored by annotated cell types from scRNA-seq analysis. D. The model embedding-generated UMAP from a GC dataset (Vikrant et al) which is a part of the training cohorts. Each dot represents a single cell. Dots are colored by annotated cell types from scRNA-seq analysis. Red circles highlighted 3 different T/NK cell sub-clusters, named as T/NK cls1-cls3. E. The UMAP of T/NK cells generated from conventional scRNA-seq analysis on the GC dataset (Vikrant et al). The T/NK subclusters identified by the embedding method were projected to the UMAP and were shown in different colors. F. GSEA dot plot shows upregulated pathways in each T/NK subcluster using GSEA analysis on GOBP database. Dot dimensions denote gene overlap counts with the GOBP database, while colors represent GSEA enrichment significance. G. The model embedding-generated UMAP from an ovarian cancer dataset (Luo et al). Each dot represents a single cell. Dots are colored by annotated cell types from scRNA-seq analysis. Red circles highlighted 2 different T/NK cell sub-clusters, named as T/NK cls1-cls2. H. The UMAP of T/NK cells generated from conventional scRNA-seq analysis on the ovarian cancer dataset (Luo et al). The T/NK subclusters identified by the embedding method were projected to the UMAP and were shown in different colors. I. Dot plot showing the expression of Tcm markers (*CCR7*, *SELL*) in the identified 2 T/NK subclusters. Scores were scaled gene expression values using gene-cell matrix from scRNA-seq data. Colors of the dots represent the expression level. The sizes of the dots represent the percentage of cells expressing the genes.

We then generated GF-based embeddings for all 1.3 million cells from 11 gastric cancer cohorts and constructed UMAPs using the same approach. Compared with UMAPs built from conventional expression-based pipelines (**Figure 1A**), GF-based embedding UMAPs revealed greater cell-type heterogeneity (**Figure 4C**). For example, in one representative cohort (Vikrant et al., 2021 [29]), GF-Llama-GC embeddings generated a UMAP partitioned by major cell types consistent with the original report; but further resolved T/NK cells into three distinct clusters (designated T/NK clusters 1–3; **Figure 4D**). When cells from these three clusters were mapped back to the expression-based UMAP using cell barcodes, they showed substantial overlap indicating that conventional pipelines could not resolve this heterogeneity (**Figure 4E**). To ensure that the three clusters were not artefacts of low cell quality, we examined standard quality-control metrics including numbers of detected genes per cell, total cell counts, and fraction of mitochondrial transcripts—and found no systematic differences among these three clusters (**Supplementary** Figure 4A). We also excluded the possibility of patient-driven batch effects (**Supplementary** Figure 4B). These controls support the three T/NK clusters representig genuine biological states. To explore their functional properties, we performed differential expression analysis followed by GSEA using both Gene Ontology Biological Process (GOBP) and Hallmark databases (**Figure 4F**; **Supplementary** Figure 4C). T/NK cluster 1 showed downregulation of immune-activation pathways and upregulation of oxidative phosphorylation, consistent with a memory-like state. Cluster 2 was enriched for immune-activation and interferon-γ response pathways, indicating an activated state. Cluster 3 was enriched for chromatin-remodelling and G2M checkpoint pathways, and exhibited low cytotoxic scores [41] based on *GZMA* and *PRF1* expression (**Supplementary** Figure 4D-E), suggesting a cycling state. Together, these results demonstrate that embeddings from GF-Llama-GC can reveal fine-grained T/NK cell heterogeneity that may not be resolved by conventional expression-based analyses.

To assess whether GF-Llama-GC can be generalised to other cancer types besides GC in cell types relevant to the tumor microenvironment, we analysed 20 ovarian cancer samples from a public scRNA-seq dataset [42]. After continual pretrained GF-Llama-GC on this dataset (see **Methods**), the model again identified distinct T/NK subgroups (**Figure 4G**) that were not detected using conventional expression-based analyses (**Figure 4H**). Quality-control metrics confirmed that the two clusters were not artefacts of low-quality cells (**Supplementary** Figure 4F), and both clusters showed similar sample distributions, ruling out patient batch effects (**Supplementary** Figure 4G). Differential expression analysis revealed that the smaller subgroup (n = 5,821) showed upregulation of central memory T-cell (Tcm) markers [43, 44], including *CCR7* (p = 2.20 × 10⁻^6^) and *SELL* (p = 7.03 × 10⁻^7^) (**Figure 4I**). These results demonstrate that embeddings from GF-Llama-GC can also resolve tumor microenviromment heterogeneity in cancers beyond GC, such as ovarian cancer, thereby highlighting the generalizability of the model across tumour contexts. In summary, GF-Llama-GC embeddings proved effective in capturing both broad and fine-grained cellular heterogeneity across a precancerous IM dataset, a GC dataset and an ovarian cancer dataset, showing the generalizability of the model across diverse tumour contexts.

### *In-silico* cellular perturbations reveal cell populations associated with disease progression

Besides revealing previously undetected cell populations, we also asked if GF-based cell embeddings can identify cell populations associated with disease progression in a robust and unbiased manner. Specifically, we adopted an analytical framework conceptually similar to *in-silico* gene perturbations using embedding shifts [1], but applied it at the cell-population level. First, we represented each sample by averaging the embeddings of all its cell sentences. Second, to simulate *in-silico* cell-type removal (KO), we computationally removed a subset of cells belonging to a given type from the sample and recalculated the sample-level embedding, evaluating if the embedding would then shift toward another state (e.g., from disease to healthy). Similarly, in a reciprocal manner, for *in-silico* cell-type transplantations we transferred cells of a given type from one sample group into another and assessed whether the recipient embedding shifted toward the donor state (see **Methods**; **Figure 5A**, **Supplementary** Figure 5A). Third, to determine thresholds of significance, the observed shifts predicted by the removal or transplantation of a given cell type were compared against shifts produced by removing or transplanting an equal number of randomly selected cells. Results were interpreted by considering both the direction of the embedding shift and statistical significance (**Figure 5A**, **Supplementary** Figure 5A).

**Figure 5.**
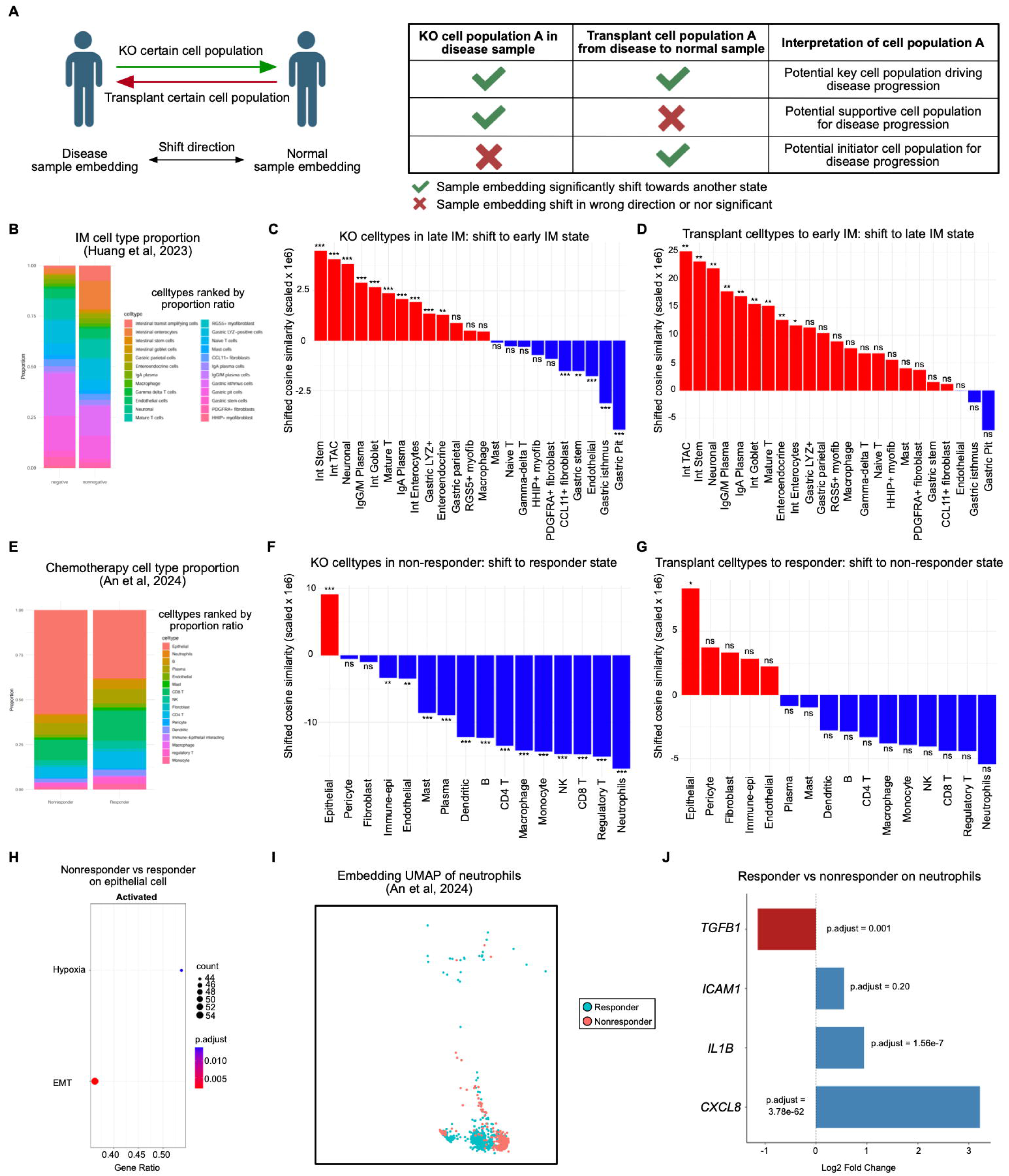
***In-silico* cell removal/transplantation using embeddings** A. Schematic depiction of performing *in-silico* cell removal/transplantation using embeddings (removal cells from disease state, transplant cells to normal state). The table (left) shows the interpretation of the result. B. The stacked bar plot showing proportions of cell types in early/ late IM samples. Bars were colored by cell types. Different cells were ranked and stacked by the proportion ratio between early/ late IM samples. C. Barplot showing the shifted cosine similarity when KO 100 cells in each cell type in late IM samples. The bars were colored by whether the shift direction is towards the early IM state (red represents the correct direction, blue represents the wrong direction). Stars represent permutation-test-derived statistical significance against KO 100 random cells for 1000 iterations (*p <0.05, ** p<0.01, *** p<0.001, ns p>=0.05). D. Barplot showing the shifted cosine similarity when transplanting 100 cells in each cell type from late to early IM samples. The bars were colored by whether the shift direction is towards the late IM state (red represents the correct direction, blue represents the wrong direction). Stars represent permutation-test-derived statistical significance against transplanting 100 random cells for 1000 iterations (*p <0.05, ** p<0.01, *** p<0.001, ns p>=0.05). E. The stacked bar plot showing proportions of cell types in chemotherapy responder/non-responder samples from a GC dataset (An et al). Bars were colored by cell types. Different cells were ranked and stacked by the proportion ratio between responder/nonresponder GC samples under chemotherapy treatment. F. Barplot showing the shifted cosine similarity when KO 100 cells in each cell type in chemotherapy non-responder samples. The bars were colored by whether the shift direction is towards the responder state (red represents the correct direction, blue represents the wrong direction). Stars represent permutation-test-derived statistical significance against KO 100 random cells for 1000 iterations (*p <0.05, ** p<0.01, *** p<0.001, ns p>=0.05). G. Barplot showing the shifted cosine similarity when transplanting 100 cells in each cell type from non-responder to responder samples. The bars were colored by whether the shift direction is towards the non-responder state (red represents the correct direction, blue represents the wrong direction). Stars represent permutation-test-derived statistical significance against transplanting 100 random cells for 1000 iterations (*p <0.05, ** p<0.01, *** p<0.001, ns p>=0.05). H. GSEA dot plot shows upregulated pathways in non-responder versus responder epithelial cells using GSEA analysis on the Hallmark database. Dot dimensions denote gene overlap counts with the Hallmark database, while colors represent GSEA enrichment significance. I. The model embedding-generated UMAP from neutrophils from samples under chemotherapy treatment (An et al). Each dot represents a single cell. Dots are colored by whether the cell is from responders or non-responders. J. Bar plot showing the log2 fold change of typical N1- and N2-neutrophil marker genes between responders and non-responders. Bars were colored by whether the gene expression was down-regulated (blue) or up-regulated (red) in non-responders.

We first applied the method to the IM dataset [23] which was not used during training and had been analysed in a previous section (**Figure 4B**). Conventional comparison of cell-type proportions between early- and late-stage IMs have highlighted Transit Amplifying Cells (TACs) as the most increased population, followed by enterocytes (**Figure 5B**). However, stem-cell-like populations are usually regarded as the key drivers of IM progression rather than these mature intestinal lineage cells [23, 45, 46]. Notably, applying the *in-silico* KO and transplantation framework, we identified intestinal stem cells as the most consistent driver population of IM progression, in line with established biology (**Figure 5C-D**). Interestingly, analysis of embedding-shift directions suggested that gastric pit cells may act as a potential “gatekeeper” population: their presence was associated with maintaining gastric identity, whereas their loss was linked to progression toward intestinal lineages (**Supplementary** Figure 5B-C). This observation is consistent with prior evidence that pit cells in IM samples can undergo metaplastic transformation and be replaced by intestinal lineage cells [23, 45]. Together, these results demonstrate that the *in-silico* KO/transplantation frameworks can be robustly applied to unseen precancer datasets, producing insights that are consistent with known biological mechanisms.

Finally, to test the cellular perturbation approach in a clinically-relevant scenario, we applied the method to a cohort of 30 GC patients who subsequently received platinum-based chemotherapy [47]. These patients were later classified as 18 responders and 12 nonresponders. Conventional cell-type proportion analysis showed that nonresponders had a higher fraction of epithelial cells, while no significant differences observed for other lineages (**Figure 5E**). Reassuringly, using our *in-silico* KO and transplantation framework, we also identified tumour epithelial cells as a key population whose removal in nonresponders or transplantation into responders consistently shifted embeddings toward the opposite clinical state (**Figure 5F-G**). Differential expression and pathway analysis showed that epithelial cells from nonresponders exhibited upregulation of EMT and hypoxia-related pathways (**Figure 5H**), consistent with prior reports linking these processes to chemotherapy resistance [48–51]. Most notably however, was that the GF-embedding framework also implicated neutrophils as another cell population linked to chemotherapy response. Specifically, *in-silico* KO of neutrophils from responder samples shifted embeddings toward the nonresponder state, while transplantation of neutrophils from responders into nonresponders partially reverted embeddings toward the responder state (**Supplementary** Figure 5D-E). Embedding-based UMAPs also separated neutrophils by response status (**Figure 5I**). Differential expression analysis showed that responder neutrophils expressed higher levels of anti-tumour N1 markers (*CXCL8*, *IL1B*, *ICAM1*), while nonresponder neutrophils upregulated the pro-tumour N2 marker *TGFB1* (**Figure 5J**) [52–54]. Together, these results demonstrate that the *in-silico* KO/transplantation framework can be applied to clinical samples, identifying both tumour epithelial cells and neutrophils as key populations linked to chemotherapy response in GC.

## Discussion

The effectiveness of AI models depends strongly on the scale and quality of training data [7]. Large language models (LLMs) have already consumed much of the available web text, and recent efforts to incorporate synthetic data have shown mixed results [55–57]. In contrast, biological data represent a vast and underexploited resource, with both real-world availability and potential clinical relevance. AI has already delivered transformative advances in biology, exemplified by AlphaFold2[58] for protein structure prediction and newer models such as Evo2 [59] for DNA sequence modelling. However, unlike proteins and DNA, which are naturally ordered sequences, RNA-seq data are typically represented as expression matrices where each entry quantifies the abundance of a gene across cells or samples, relative to other genes. As gene expression matrices lack intrinsic order (ie genes that are largely unrelated may situated adjacent to one another simply due to expression values), they do not map naturally onto sequence-based deep learning architectures such as transformers. While several groups have begun to explore single-cell foundation models, most existing approaches still rely on ranked-gene encodings or hybrid schemes that combine ranks with expression values, and innovations in how gene expression is represented remains limited. Thus, to achieve an RNA-seq model with impact comparable to AlphaFold2 in structural biology, new encoding strategies for expression matrices are urgently needed.

In this study, Gene-family (GF) representations were developed to address limitations of ranked-gene (RG) encoding, by grouping genes into families that reflect biological functions and pathways. This approach provides a more interpretable structure than simply ordering genes by expression level. To evaluate the impact of GF encoding, we focused on benchmarks designed to test whether models capture biologically meaningful relationships, rather than tasks such as cell annotation or sample integration that require fine-tuning on the test data and may not directly reflect the quality of the pre-trained embeddings. Specifically, we assessed whether models could recover relationships among genes within the same cell type and within known pathways or gene programs. Compared with RG encoding, GF-encoded training yielded higher readability scores, faster convergence during domain-adaptive pre-training, and superior performance in distinguishing related from unrelated genes at both the individual-gene and gene-program levels. GF-Llama, in particular, accurately recovered known gene–gene relationships, produced coherent cell type-specific gene association networks, and achieved the highest separation scores for both predefined gene families and independent pathway gene sets. These gains were reproducible across backbones and extended to gene programs not seen during training, underscoring the generalisability of this GF approach.

GF encoding also showed superior performance on generative evaluations, ranging from single-token prediction to whole-cell sentence generation. Moreover, using GF-Llama-GC, we generated complete cell sentences which reproduced expected marker profiles under each condition, showing the potential in stimulating synthetic cell sentences in customized conditions. Our results also underscore that backbone choice and tokenizer design substantially influence model performance, with Llama3.1 outperforming Qwen3 despite lower general-purpose leaderboard rankings, likely due to more efficient tokenisation of gene symbols. Together, these findings highlight GF encoding and careful backbone–tokenizer selection as key factors for building high-performing, domain-adapted generative models in single-cell biology.

Most previously developed single-cell foundation models have focused on applications that substantially overlap with established bioinformatics pipelines, limiting their novelty. In this study, we sought to address fundamental challenges in scRNA-seq by introducing two distinct applications of GF-Llama-GC: (1) identification of cellular heterogeneity and (2) identification of key cell populations associated with disease states. Using embedding-based UMAPs, GF-Llama-GC robustly recovered gastric and intestinal epithelial lineages in a zero-shot precancerous dataset, revealed novel T/NK subclusters with distinct functional programs in GC, and generalised to ovarian cancer where they uncovered Tcm populations missed by conventional pipelines. These results highlight the ability of gene-family–based foundation models not only to reproduce known biology but also to discover previously hidden cellular states, underscoring their potential as a powerful framework for cross-and pan-cancer single-cell analysis.

In parallel, our *in-silico* cell removal and transplantation conceptual framework provides an unbiased means of identifying cell types that influence sample-level states across populations. In an IM dataset, although intestinal stem cells are not a major cell type in either the cell number or the cell proportion between early- and late-stage IM samples, the method still predicted intestinal stem cells as the key cell type associated with IM progression, in line with biological facts. In a chemotherapy cohort, besides the tumor epithelial cells, the method also highlighted a potentially important role for neutrophil subtypes in shaping therapeutic response, consistent with recent literature [60, 61]. These methods are broadly applicable in clinical settings. For example, they can be used to dissect cellular heterogeneity and state-driving populations that distinguish immunotherapy responders from non-responders, early-from late-stage cancers, or intestinal-from diffuse-type gastric cancers. Furthermore, their generalizability extends beyond gastric cancer, as demonstrated by the identification of T-cell heterogeneity in ovarian peritoneal metastases.

This study should be considered as a proof-of-concept report and has several limitations. First, although we defined 29 gene families intended to capture major aspects of cellular activity, some functional categories are inevitably missing. In the current framework, genes outside the predefined families were grouped into an “Other genes” category. Future refinements of gene-family definitions will be definitely required. For example, model attention patterns could be used to explore associations between genes and established families, potentially enabling data-driven expansion or redefinition of gene families in future work. Second, our evaluation was restricted to GC datasets and two open-source backbone models. While sufficient to demonstrate proof-of-concept, broader testing will be required. Because GF encoding functions as a data representation strategy, it can in principle be applied to other disease settings and to more powerful foundation models. Applying GF encoding to well-curated pan-cancer datasets and training on state-of-the-art LLMs with larger parameter scales could substantially improve model performance. Such efforts may eventually enable advances in scRNA-seq analysis comparable in impact to breakthroughs such as AlphaFold2 in protein structure prediction.

## Methods

### Development of the GC-atlas scRNA-seq corpus

We compiled single-cell expression data and matched clinical metadata from 11 public gastric cancer cohorts (**Supplementary Table 1**). For cohorts which provide raw FASTQ data, these data were aligned to GRCh38 and generated cell-gene count matrix using CellRanger v7.0. Metadata fields used included relevant information such as Accession number, patient ID, sample ID, tumor/normal, age, gender, MSI, Lauren subtype, H. pylori status, stage and so on (**Supplementary Table 1**). Gene symbols were standardised to HGNC. Using Scanpy v1.11, we merged all cells using scanpy.concat(join=“outer”), obtaining 1,355,628 cells and 40,596 genes. We computed per-cell QC metrics (total_counts, n_genes_by_counts, pct_counts_mt, with MT genes defined by the “MT-” prefix). Cells were excluded if total UMIs < 600 or > 50,000, genes detected < 300, or pct_counts_mt > 50%. Genes expressed in < 3 cells in each dataset were removed. After this step, 1,315,032 cells and 36,017 genes remained. Potential doublets were flagged using Scrublet (implemented under Scanpy v1.11) using default settings, yielding 1,313,051 cells and 36,017 genes for downstream analysis. We next ran scVI v1.3, used raw counts and recorded samples as the batch variable. We selected 6,000 HVGs with pp.highly_variable_genes(flavor=’seurat_v3’, batch_key=’sample’). We trained scVI with parameters: n_layers=2, n_latent=30, gene_likelihood=’nb’ on 1*A100 GPUs. Batch covariates were modelled via samples. We then computed a kNN graph on the scVI latent space, followed by UMAP (min_dist=0.3) and Leiden clustering. Differential expression was performed with tl.rank_genes_groups using the Wilcoxon rank-sum test, comparing each cluster vs all other cells, with Benjamini–Hochberg FDR control. We then annotated the cell type for each cluster manually based on the top differentially expressed genes.

### Gene family encoding to generate cell sentences

We defined 29 gene families to cover general cellular functions (15 families) and tumour-specific processes (14 families) using curated resources including Gene Ontology (GO), Reactome, DepMap, OncoKB, and primary literature (**Supplementary Table 2**). By design, families include genes with both activating and inhibitory roles (e.g., stimulatory and inhibitory checkpoint genes are grouped within Immune state), i.e., no polarity is encoded at the family level. After de-duplication across sources, the catalogue covers 13,647 unique genes. An additional 30th family, “Other genes,” contains genes not included in the above 29 families. A gene may belong to multiple families when supported by source annotations; duplicates are retained at the catalogue level. To visualise relationships among families, we computed the Jaccard index for all family pairs and built a family network in R v4.1 using igraph v1.2, tidygraph v1.2, and ggraph v2.0. Edge width encodes Jaccard similarity; node size encodes the number of genes per family; node colour denotes general vs tumour-specific families.

Each cell sentence begins with a metadata sub-sentence generated from available fields using the following template: *“The {cell_type} cell originates from {tumour/normal} {tissue} in a gastric cancer patient. Patient characteristics — gender: {gender}; age: {age}; Lauren subtype: {lauren}; H. pylori status: {hpylori}; stage: {stage}; location: {location}; TCGA subtype: {tcga}; EBV status: {ebv}; HER2 status: {her2}.”*

Fields missing for a cell are omitted (no placeholder token). For each cell, we partitioned non-zero expressed genes (count > 0) into 30 subgroups: the 29 predefined families plus Other genes. Within each subgroup, genes were ranked by per-cell expression (raw count). To balance information content with model context length, we retained the top 50 genes per predefined family and the top 500 genes in Other genes (if fewer than the cut-off exist, all available genes were used). If a gene belongs to multiple families, it is listed once per family but never duplicated within a family. We realised sub-sentences using the following formats:

- Family sub-sentence: *“The top-ranked expressed genes in {family_name} are [{GENE1} {GENE2}, …].”*
- Other genes sub-sentence: *“The top-ranked remaining genes are [{GENE1}*

*{GENE2} …].”*

A complete GF-encoded cell sentence concatenates the metadata sub-sentence followed by the family sub-sentences in a fixed order defined in **Supplementary Table 2**. For benchmarking, we also generated 100,000 RG-encoded sentences, which keep the same metadata but list a single ranked gene list:

*“The top-ranked genes are [{GENE1} {GENE2} …].”*

All cell sentences were saved in JSON Lines (JSONL) with fields { “text”: {Cell Sentence}}. Files were shuffled at the cell level.

### Adaptive-pretraining on public LLM models

We used two open-source backbones: Llama3.1-8B-Base (https://huggingface.co/meta-llama/Llama-3.1-8B) and Qwen3-8B-Base (https://huggingface.co/Qwen/Qwen3-8B). Unless stated otherwise, we used each backbone’s native tokenizer. Continual pre-training on cell sentences was performed with Axolotl v0.9, which wraps PyTorch v2.7 and Transformers v4.51. The objective was causal language modelling (next-token cross-entropy). We used a maximum sequence length of 4096 tokens and packing = true. Training was on full parameters. Key Axolotl YAML settings were: optimizer = AdamW, learning rate = 2e-5, learning rate scheduler = cosine, warmup = 0.01, weight decay = 0.0, mixed precision = bf16. Distributed training used DeepSpeed ZeRO-3. We trained for 1 epoch on 100,000 cell sentences (GF and RG) for benchmarking, and for 3 epochs on 1.3 M GF-encoded sentences for the final model (GF-Llama-GC). Full YAMLs are provided in GitHub (will be available after paper acceptance). Training ran on 8× NVIDIA H200 (80 GB) GPUs on the A*CRC Singapore cluster.

### Readability score of cell sentences

We assessed the readability of GF- and RG-encoded cell sentences using an external LLM judge as a proxy metric. Specifically, we queried Gemini-2.0-flash with a fixed prompt instructing the model to rate how understandable a sentence would be to a cancer cell-biology expert on a 1–10 scale and to return only a numeric score. We evaluated 100 randomly sampled sentences per encoding (GF and RG). Metadata templates were identical across encodings. The judge received only the sentence string (no encoding label). The exact prompt was:

*“You are a biomedical language expert. I will give you a sentence that contains many gene names and technical terms. Your task is to evaluate how understandable this sentence would be to a cancer cell biology expert reader — assuming they are familiar with gene markers, pathways and their functions. Rate the sentence’s understandability on a scale from 1 to 10, where 1 means confusing or unreadable, and 10 means very easy to understand for an expert. Only return the numeric score.”*

### Bio-knowledge-based benchmarking using embeddings

We benchmarked four models trained in this study (GF-Llama, RG-Llama, GF-Qwen, RG-Qwen) against three baselines: Llama3.1-8B-Base (https://huggingface.co/meta-llama/Llama-3.1-8B), Qwen3-8B Base (https://huggingface.co/Qwen/Qwen3-8B), and Geneformer-V2-104M_CLcancer (https://huggingface.co/ctheodoris/Geneformer/). For Llama and Qwen models, to obtain a gene-level vector from decoder-only backbones, input genes were tokenised with the native tokenizer of each backbone. For genes that split into multiple subword tokens, we computed the mean of the final-layer hidden states across all subword positions corresponding to the gene token span. The resulting vectors were 4096-dimensional (Llama 3.1 8B, Qwen 3 8B). Embeddings were directly used for cosine similarity calculation in similarity analyses. Geneformer provides a gene-token vocabulary with a 1:1 mapping between tokens and genes. We retrieved the token embedding/last hidden state for each gene by passing the corresponding gene token. The resulting vectors were 512-dimensional. All between-model comparisons were performed within each model’s embedding space (i.e., we did not compute cross-model distances). Where cross-model summaries were shown, they report model-specific metrics (e.g., median within-set cosine similarity) rather than raw vector norms. We used the gene-level embeddings (see above) to assess whether models recover expected biological relationships among cell-type marker sets curated in **Supplementary Table 2**. For each model, pairwise cosine similarities between gene vectors were computed within and between marker sets. We constructed gene–gene KNN graphs with k = 10 (NearestNeighbors(metric=’cosine’, algorithm=’auto’, n_neighbors=10)). Graphs were symmetrised by keeping mutual-KNN edges. For visualisation, we retained the top 50 edges by cosine similarity per graph to reduce clutter. We rendered embedding-based gene association networks in igraph v1.2; edge width was proportional to similarity (min–max scaled within the graph), and node colour reflected the canonical marker lineage from **Supplementary Table 2**. As a positive control, we examined RPL genes (housekeeping, functionally homogeneous). As a negative control, we sampled pairs across distinct immune programmes (inhibitory checkpoints, stimulatory checkpoints, chemokines, cytokines, MHC) (gene lists and sources in **Supplementary Table 2**). For each model, we generated iteration-level means of pairwise cosine similarities for each group (positive, negative, null) via repeated random sampling (n_pairs = 100; n_iter = 100). We summarised distributions with density plots and descriptive statistics, reported the 95% central interval of the null distribution, and estimated the common-language probability P(X>Y) between groups.

We evaluated whether gene embeddings separate functionally distinct pathways/gene programs. We analysed Gene Family 7, Gene Family 18, Hallmark EMT, Hallmark IFN-γ. Genes annotated to multiple sets were still kept and annotated as “mixed”. For each model, we compute cosine similarities of gene vectors. For visualization only, embedding matrices were z-scored per dimension across genes, followed by PCA (n_components=20) and UMAP. UMAPs were generated separately for each model and gene set combination. We also quantified separation using cosine similarity on the original embeddings. We defined the normalized intra/inter cosine similarity R value as follows:

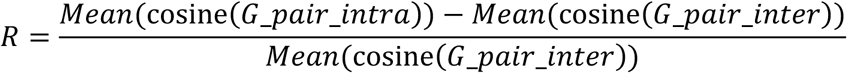

Where cosine(G_pair_intra) denotes cosine similarity between embeddings from all unique combinations of two genes that belong to the same gene program, cosine(G_pair_inter) denotes cosine similarity between embeddings from all unique combinations of two genes that belong to distinct gene programs. We need to note that for most models, Mean(cosine(G_pair_inter)) is a value that is very close to 1.

### Generative-based benchmarking

We designed a cell prompt consists of consisting of (i) a metadata sentence and (ii) two cell-type marker genes as hints. The same prompt set was used for all models. Using randomly generated cell prompts from different cells, we calculated the likelihood of the top-1 probability and top-5 cumulative probability mass over the full vocabulary when we input the cell prompts to the model with deterministic decoding (do_sample=False, temperature=0, top_p=1). For whole cell generation, we first generated the same cell prompts (2 hint genes from cell type) from 100 cell sentences, and let the model continue generating at most 1500 tokens, then compared the generated cell sentences with the reference cell sentences using BERTScore from the bert-score package v0.3 in Python. To assess conditioning on metadata alone, we also constructed 100 prompts without hint genes (metadata only) and repeated the generation and BERTScore evaluation with identical decoding settings.

For metadata scenarios in **Figure 3C–E** (tumour vs adjacent normal; stage I vs stage IV; Lauren intestinal vs diffuse), we used the same prompt template with condition fields set accordingly and generated up to 6,000 tokens (max_new_tokens=6000, do_sample=False). We compared condition-specific marker profiles, highlighted a priori expected markers supported by literature.

### Processing of validation scRNA-seq dataset

We used a public intestinal metaplasia (IM) scRNA-seq dataset [23] comprising 71933 cells from 18 patients. Cells and matched clinical fields were converted to GF-encoded cell sentences using the procedure described above. For IM we used a shortened metadata template:

*“The {cell_type} cell originates from {tumour/normal} gastric tissue in a patient.”*

The ovarian peritoneal metastasis dataset (Luo et al., [42]) contained 209,117 cells from 20 samples; its metadata template was:

*“The {cell_type} cell originates from {tumour/normal} tissue in an ovarian metastasis patient.”*

For GC datasets from Vikrant et al [29] and An et al [62], these GC cohorts were part of the training corpus. We therefore reused their training sentences to compute embeddings for visualisation only; results from these cohorts were not interpreted as independent evidence of model generalisation.

All sentences were tokenised with the default backbone Llama tokenizer. We note that the IM dataset was not used for training or fine-tuning. We directly encoded sentences with GF-Llama-GC to obtain embeddings for downstream analyses. Because ovarian cancer differs from GC, we performed one epoch of domain-adaptive pre-training on the ovarian sentences using Axolotl with the same objective (causal LM) and same hyperparameters as in GC training. The adapted checkpoint was then used to embed the ovarian sentences as above.

Similar to gene-level embedding processing and visualization, for all cell embeddings from various datasets, we first scaled the embeddings using Z-transformation, then performed PCA (dims = 30), and finally visualised the data in UMAP space.

### *In-silico* cell removal/transplantation

We define two patient groups-population A (e.g., healthy samples) and population B (e.g., disease samples)-each comprising a set of patients with cell-level embeddings derived as described above (token pooling from final layer). For in-silico cell removal in population A, we first calculated the original cell embeddings EA from the cell sentences. Next, for every annotated cell type in the dataset, we randomly remove 100 cells and re-generated the cell embeddings E’A_celltype_KO, respectively. If the cell number for a certain cell type is less than 100, we will remove all cells in this cell type. For each E’A_celltype_KO from different cell types, we measured the changes in the difference of cosine similarity among populations using the following equation:

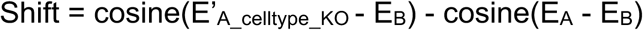

To measure the significance of the shift, we performed a permutation test which removed 100 randomly selected cells from population A, and repeated for 1,000 iterations to get the background differences.

Similarly, for in-silico cell transplantation, we randomly added 100 cells from each cell type in population B to population A, and measured the shift distance using the following equation:

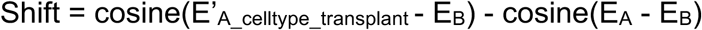

## Data and code availability

The four benchmarking checkpoints (GF-Llama, RG-Llama, GF-Qwen, RG-Qwen) and the scaled model GF-Llama-GC are deposited on Hugging Face with private reviewer links provided at submission and will be publicly available on publication. The 1.3 million GF cell sentences are released as a dataset on Hugging Face Datasets. We do not re-host raw scRNA-seq matrices and instead supply a manifest with accessions for all source cohorts in **Supplementary Table 1**. In-house code for sentence generation, training (Axolotl YAMLs), embedding extraction, and *in-silico* cell removal/transplantation is on GitHub (publicly available on publication). Additional information is available from the corresponding author on reasonable request.

## Grant support and Acknowledgements

This work was supported by the National Medical Research Council Singapore Translational Research Investigator (STaR) award (MOH-000967). This research was also supported by the Duke-NUS Core funding. This work was supported by the A*STAR Computational Resource Centre (A*CRC) through the use of its high performance computing facilities.

## Supplementary Figure Legends

**Supplementary Figure 1.**
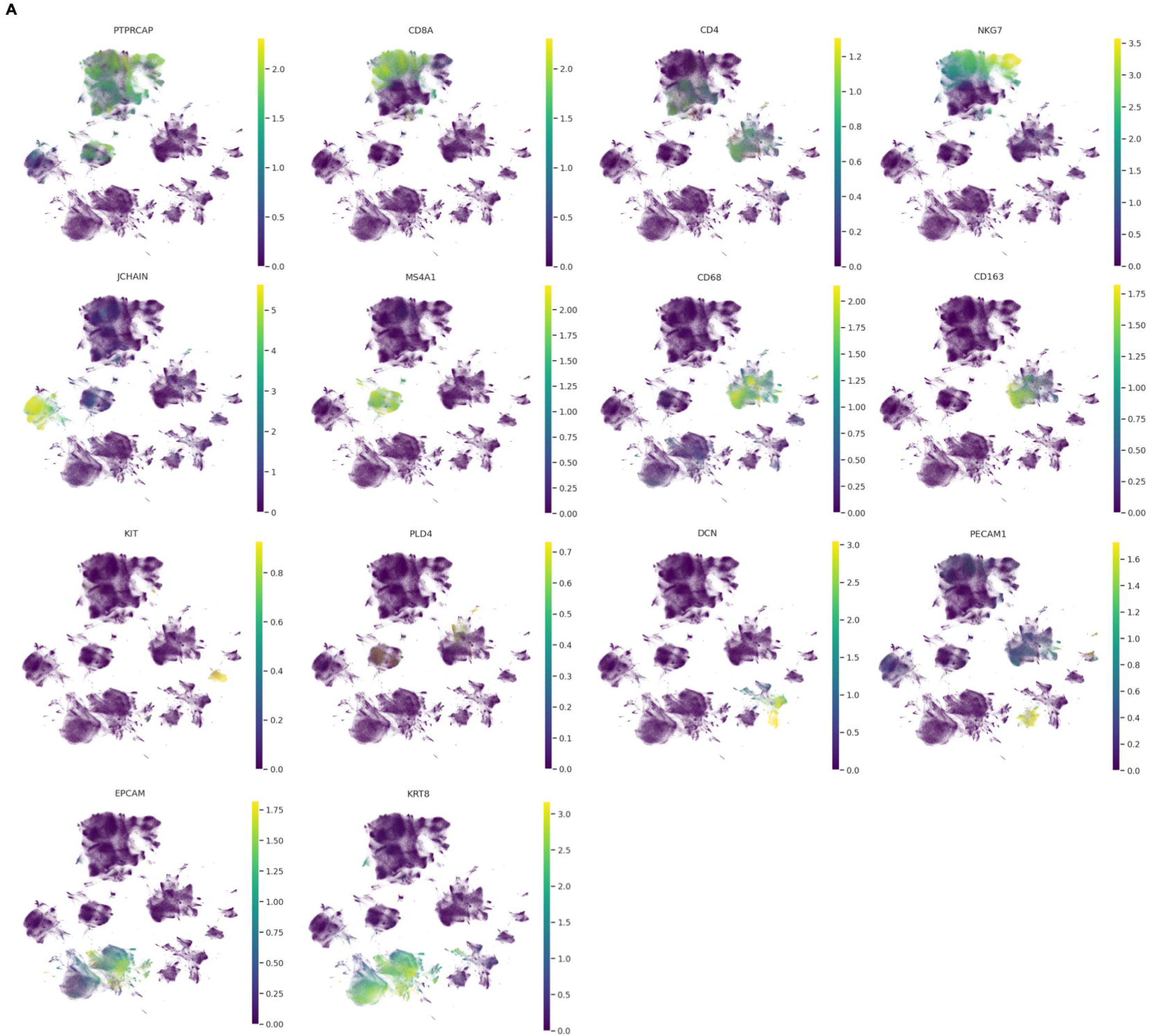
A. Dim plot of cell-type-specific markers on integrated GC scRNA-seq data (1.3 million cells). The values represent normalized expression data. Markers include: *PTPRCPR, CD8A, CD4, NKG7, JCHAIN, MS4A1, CD68, CD163, KIT, PLD4, DCN, PECAM1, EPCAM, KRT8*.

**Supplementary Figure 2.**
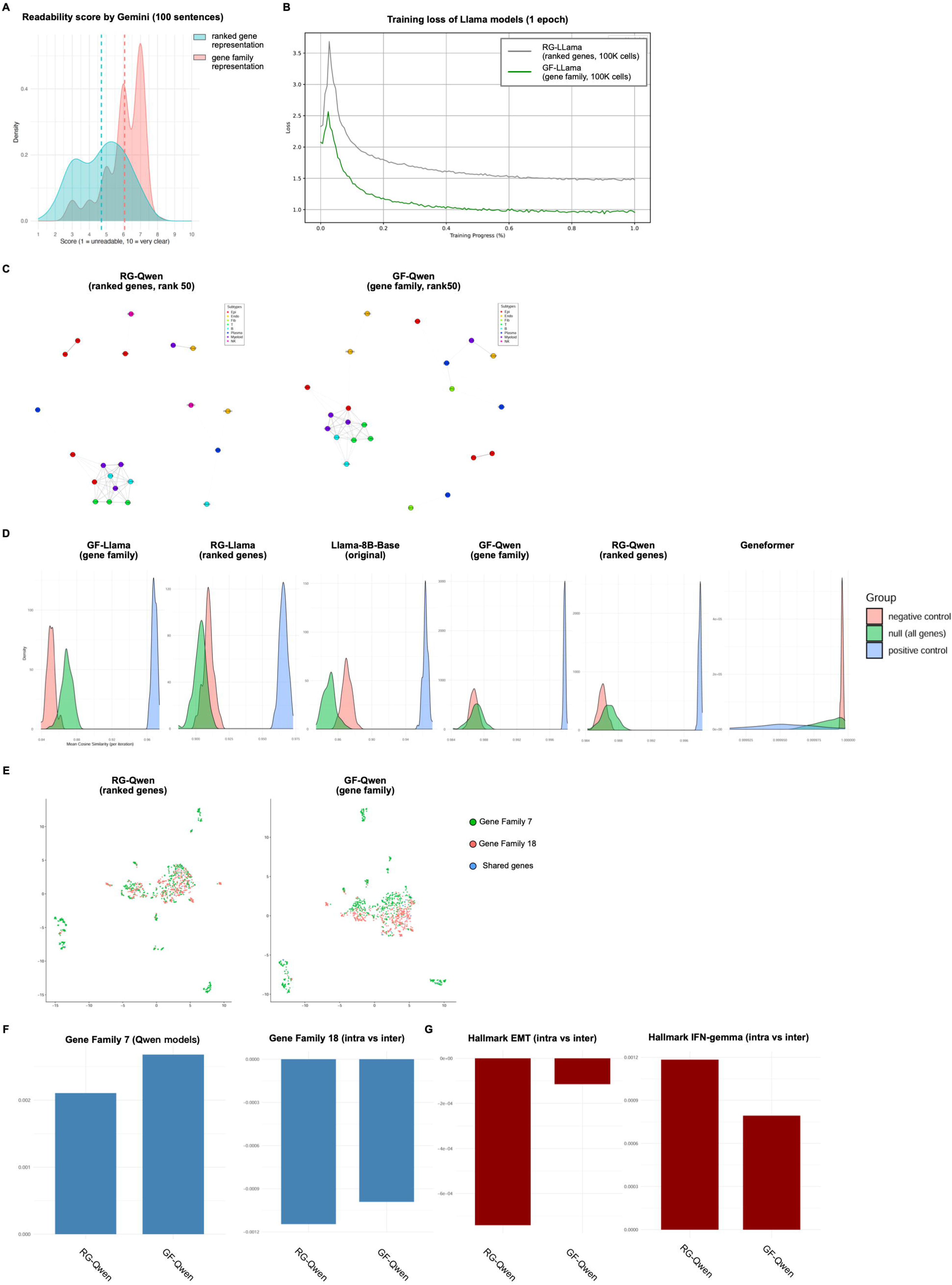
A. Density curve plot of the readability scores defined by Gemini-2.5-Flash. Blue curves represent scores for RG-encoded cell sentences, and red curves represent scores for GF-encoded cell sentences. Dashed lines represent median scores. B. The training loss curves for Llama models (GF-Llama and RG-Llama). The grey line represents the training loss for RG-Llama, while the green line represents the training loss for GF-Llama. Each model was trained on 100K cells for 1 epoch. C. Gene correlation network plot defined using KNN and cosine similarities of cell-type-specific gene embeddings for Qwen models. Each dot represents a gene, and the dot was colored by cell types. The link between two dots represents that these two genes have a high cosine similarity ranked top 50 among all gene pairs. D. Density plot of the cosine similarity between genes in the positive control group (colored in blue), the negative control group (colored in read), and the background random gene group (colored in green). E. UMAP projections of gene embeddings from a mixture of defined Gene Family 7 (protein translation-related genes) and Gene Family 18 (oncogene-related genes) for Qwen models. Each dot represents a gene, and the dot was colored by different gene families. F. Barplots showing the defined normalized intra/inter cosine similarity scores on Gene Family 7, Gene Family 18 and 2 unseen Hallmark pathways (EMT and IFN-γ) that are not defined in gene families in Qwen models. Positive values show the model can distinguish among different gene programs/pathways, and the magnitude of the values represents how well it can separate gene programs/pathways.

**Supplementary Figure 3.**
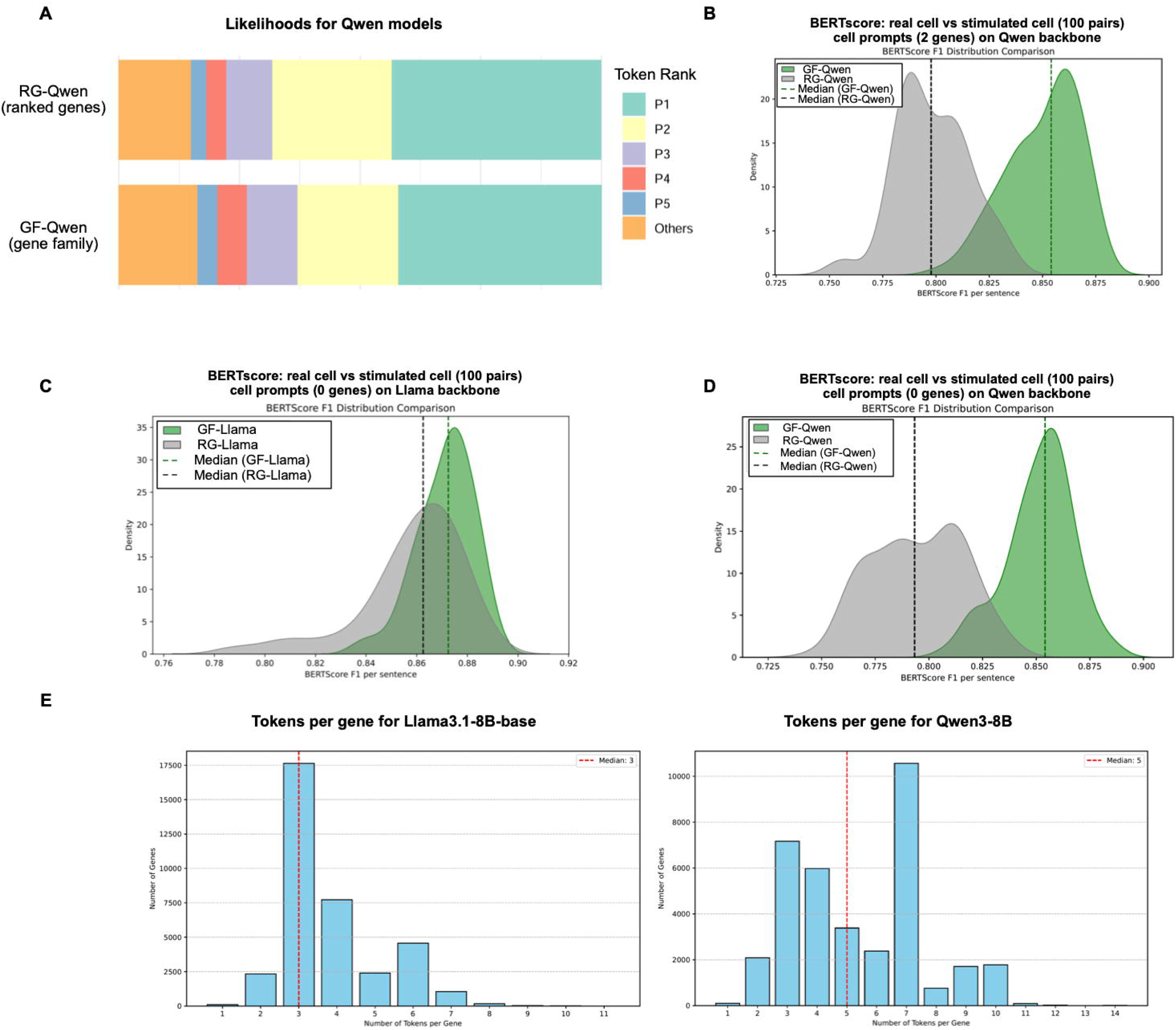
A. Bar plots showing the distribution of the likelihood of the next predicted token from GF-Qwen and RG-Qwen. The likelihood of the top 5 candidate tokens and the rest was colored. The result takes the mean value from 1,000 sentences. B. Density plot of the BERTscore F1 values for GF-Qwen and RG-Qwen between 100 pairs of stimulated cell sentences (with 2 hint genes) and reference sentences. The dotted line indicates the median value of each model. Colors represent the F1 values from different models. P-value is 7.91e^-13^ by the two-sided Mann–Whitney U test. C. Density plot of the BERTscore F1 values for GF-Llama and RG-Llama between 100 pairs of stimulated cell sentences (with 0 hint genes) and reference sentences. The dotted line indicates the median value of each model. Colors represent the F1 values from different models. P-value is 7.91e^-13^ by the two-sided Mann–Whitney U test. D. Density plot of the BERTscore F1 values for GF-Qwen and RG-Qwen between 100 pairs of stimulated cell sentences (with 0 hint genes) and reference sentences. The dotted line indicates the median value of each model. Colors represent the F1 values from different models. P-value is 7.91e^-13^ by the two-sided Mann–Whitney U test. E. Bar plot of the frequency distribution of the number of tokens per gene symbol using tokenizers from Llama3.1-8B-base (left) and Qwen3-8B (right). Red dotted lines represent median numbers of tokens per gene symbol.

**Supplementary Figure 4.**
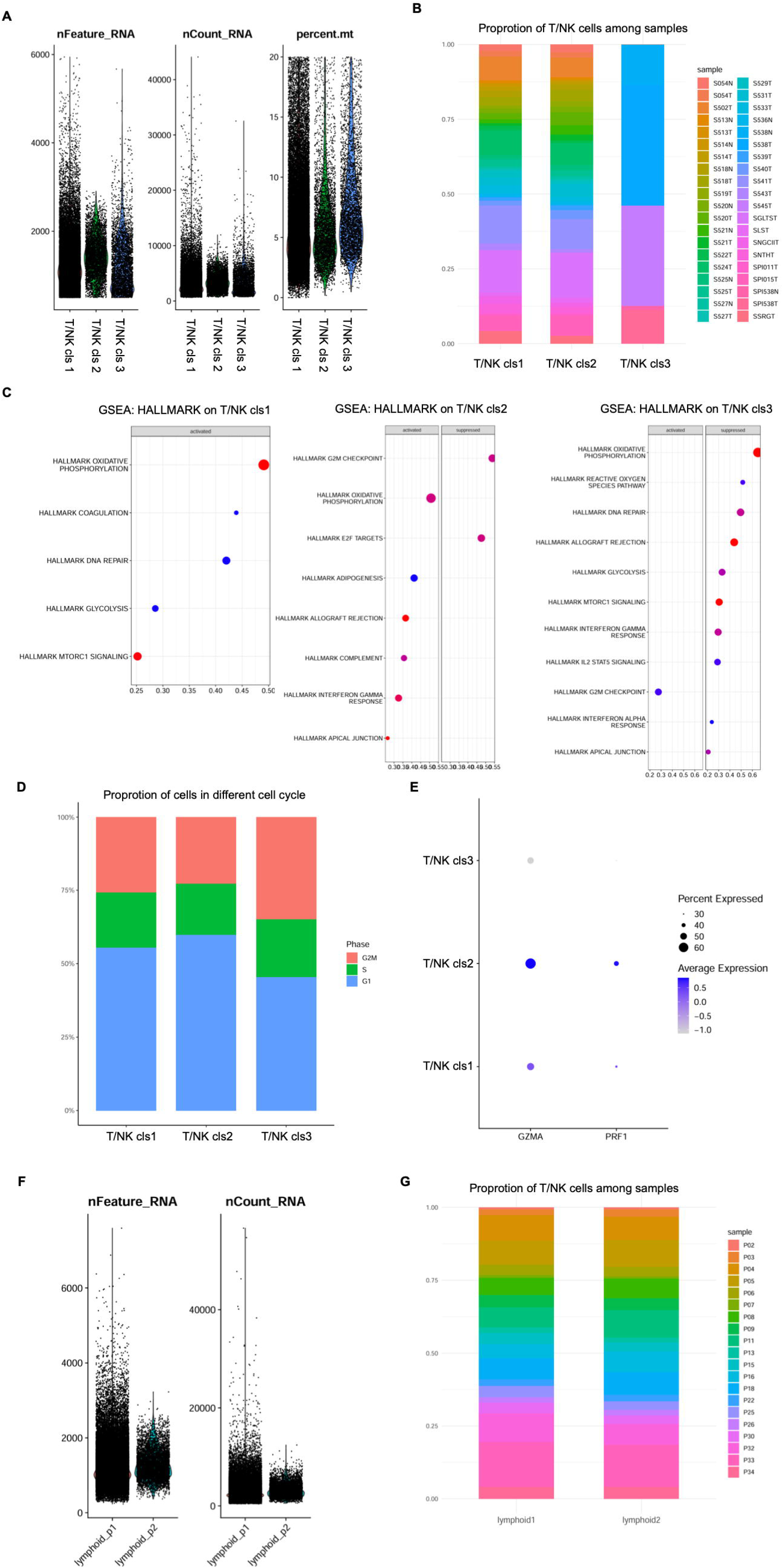
A. Violin plot showing the quality control metrics of the 3 T/NK subcluster cells in a GC dataset (Vikrant et al). Each dot represents a cell. B. The stacked bar plot showing sample proportions in the 3 T/NK subclusters. Bars were colored by samples. C. GSEA dot plot shows upregulated pathways in each T/NK subgroup using GSEA analysis on the Hallmark database. Dot dimensions denote gene overlap counts with the Hallmark database, while colors represent GSEA enrichment significance. D. Bar plots of cells in different cell cycles (G2M, S, G1) in each T/NK subgroup. Bars are colored by different cell cycles. E. Dot plot showing the expression of cytotoxic score markers (*GZMA*, *PRF1*) in the identified 3 T/NK subclusters. Scores were scaled gene expression values using gene-cell matrix from scRNA-seq data. Colors of the dots represent the expression level. The sizes of the dots represent the percentage of cells expressing the genes. F. Violin plot showing the quality control metrics of the 2 T/NK subcluster cells in an ovarian cancer dataset (Luo et al). Each dot represents a cell. G. The stacked bar plot showing sample proportions in the 2 T/NK subclusters. Bars were colored by samples.

**Supplementary Figure 5.**
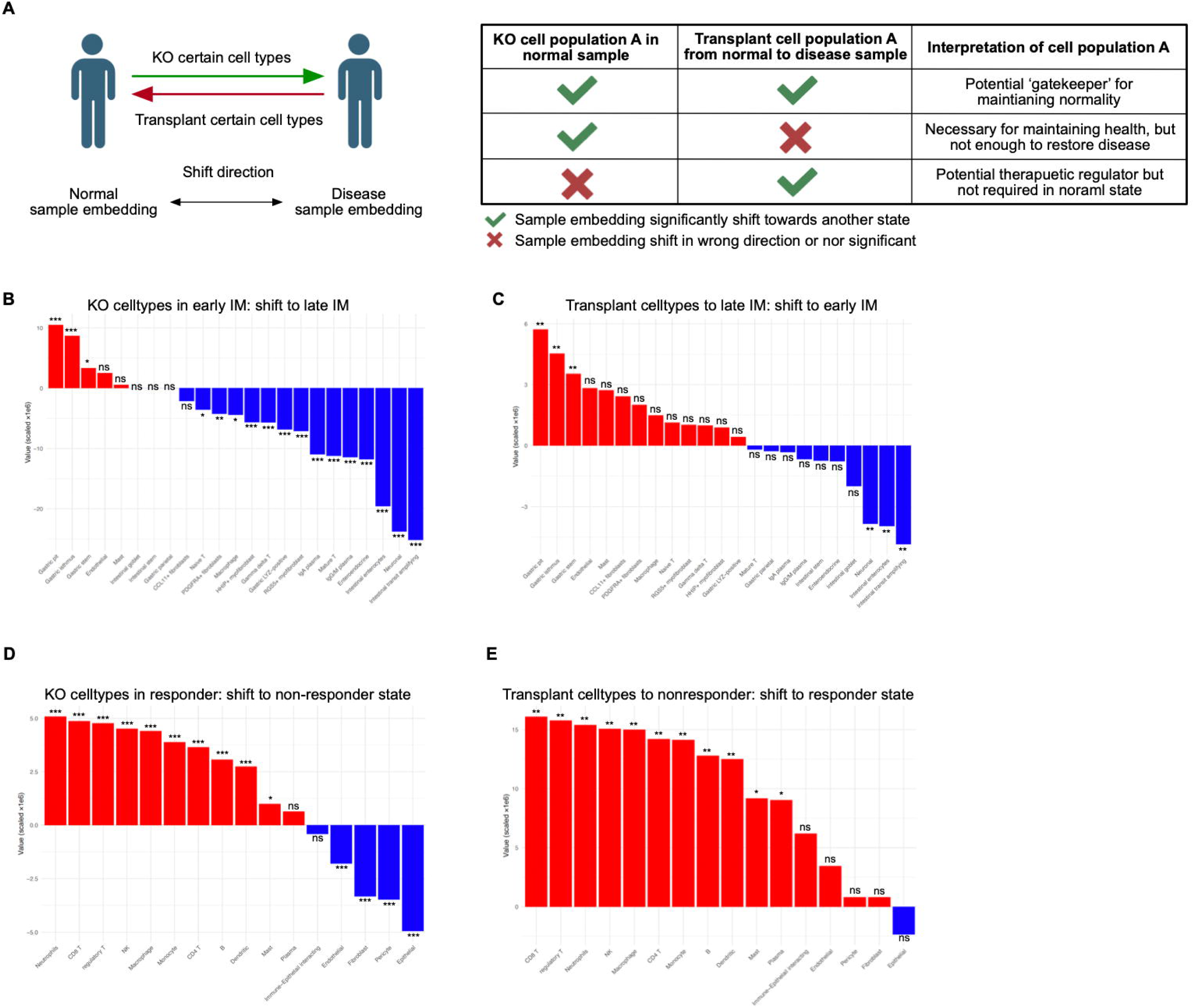
A. Schematic depiction of performing *in-silico* cell removal/transplantation using embeddings from another side (removal cells from normal state, transplant cells to disease state). The table (left) shows the interpretation of the result. B. Barplot showing the shifted cosine similarity when KO 100 cells in each cell type in early IM samples. The bars were colored by whether the shift direction is towards the late IM state (red represents the correct direction, blue represents the wrong direction). Stars represent permutation-test-derived statistical significance against KO 100 random cells for 1000 iterations (*p <0.05, ** p<0.01, *** p<0.001, ns p>=0.05). C. Barplot showing the shifted cosine similarity when transplanting 100 cells in each cell type from early to late IM samples. The bars were colored by whether the shift direction is towards the early IM state (red represents the correct direction, blue represents the wrong direction). Stars represent permutation-test-derived statistical significance against transplanting 100 random cells for 1000 iterations (*p <0.05, ** p<0.01, *** p<0.001, ns p>=0.05). D. Barplot showing the shifted cosine similarity when KO 100 cells in each cell type in chemotherapy responder samples. The bars were colored by whether the shift direction is towards the non-responder state (red represents the correct direction, blue represents the wrong direction). Stars represent permutation-test-derived statistical significance against KO 100 random cells for 1000 iterations (*p <0.05, ** p<0.01, *** p<0.001, ns p>=0.05). E. Barplot showing the shifted cosine similarity when transplanting 100 cells in each cell type from responder to non-responder samples. The bars were colored by whether the shift direction is towards the responder state (red represents the correct direction, blue represents the wrong direction). Stars represent permutation-test-derived statistical significance against transplanting 100 random cells for 1000 iterations (*p <0.05, ** p<0.01, *** p<0.001, ns p>=0.05).

